# Intracellular pH links energy metabolism to lymphocyte death and proliferation

**DOI:** 10.1101/2021.10.29.466539

**Authors:** Wei-ping Zeng, Shuangshuang Yang, Baohua Zhou

## Abstract

The role of intracellular pH (pHi) of lymphocytes in the control of the magnitude of immune response is unknown. We found that low pHi induces apoptosis of proliferating lymphocytes, whereas high pHi promotes their survival. In the ovalbumin sensitization and challenge model, energy metabolism is a major mechanism for regulating pHi. TCA cycle using carbohydrates lowers whereas glutaminolysis or aerobic glycolysis increases pHi. Proliferation powered by high mitochondrial membrane potentials (MMPs) in lymphocytes of low but not high pHi causes apoptosis. After antigenic challenge, lymphocytes of high pHi gradually increase and assume a positive relation between pHi and MMPs while lymphocytes of low pHi and an inverse relation between pHi and MMPs diminish. This change is at least partly dependent on glutaminolysis and aerobic glycolysis.

**One Sentence Summary:** TCA cycle fueled with carbohydrates but not glutaminolysis or aerobic glycolysis lowers intracellular pH to induce apoptosis.

## INTRODUCTIOIN

Cell death and proliferation play fundamental roles in many biological processes including immune response. A typical in vivo immune response peaks at around one week and may last for several weeks, during which lymphocytes expand and execute their effector functions (*1, 2*). Eventually most of the expanded lymphocytes die of apoptosis (*3, 4*). During this dynamic process, continuous rebalance of cell proliferation and apoptosis determines the magnitude of the immune response. An early study suggests that selective use of energy metabolic pathways may play a role in determining the magnitude of immune response. It was found that the increase of energy production via aerobic glycolysis strongly correlates with lymphocyte proliferation (*5*). This phenomenon was consistent with the previously described preferential use of aerobic glycolysis over mitochondrial respiration for energy production in tumor cells and other proliferating normal cells, which has come to be known as the Warburg effect (*6–10*). It is estimated that 70-80% of human cancers use the Warburg effect for energy production (*11*), and the contribution of aerobic glycolysis to total energy production ranges from less than 5% to more than 50% (*12*). The relative contribution of aerobic glycolysis to total energy production in lymphocytes is less clear.

In more recent years, studies have identified the PI3K/AKT/mTOR pathway as the mechanism for promoting the Warburg effect. This pathway directly or through HIF-1α regulates glucose import to the cell and the expression or activities of metabolic enzymes that favor glycolysis over mitochondrial carbon oxidation (*8, 13, 14*). Studies have also unveiled an unexpected role of energy metabolism in T cell subset differentiation. It is found that the aerobic glycolysis pathway is required for optimal T helper effector cell differentiation, whereas its inhibition enhances the differentiation of Treg cells and memory CD8 T cells (*15–22*). The suppression of memory CD8 T cells formation by aerobic glycolysis is caused by AKT-mediated phosphorylation of Foxo1 using the glycolytic ATP as substrate (*22*). On the other hand, mitochondrial respiration fueled by fatty acid β-oxidation promotes Treg and memory CD8 T cell differentiation (*18, 19*). Blocking aerobic glycolysis pathway or glutaminolysis, which is another energy metabolic feature of proliferating cells, have also been shown to impair T cell proliferation (*16, 17, 21, 22*), the precise mechanism for which was not investigated but under the current framework it is presumably due to insufficient biomass and ATP synthesis.

The end product of the Warburg effect lactate is exported out of the cell so that it creates an acidic extracellular environment. Many studies have investigated the biological consequences of the extracellular acidosis. In tumors, the acidic extracellular microenvironment facilitates tumor cell invasion of local tissues (*23, 24*). Extracellular acidosis is also a long recognized feature of inflammation (*25*). In general, extracellular acidosis dampens the effector functions and anti-tumor activities of innate immune cells such as the neutrophils, tumor-associated macrophages, and NK cells (*26–29*). It also suppresses T cell-mediated immunity by inhibiting cytotoxicity and blocking cell cycle progression and the expression of IL-2 and IFN-γ (*30*).

Another salient feature of the Warburg effect is its low efficiency of energy production as it net produces only 2 ATPs per glucose as opposed to 34 ATPs through mitochondrial respiration (*31*). Intriguingly, glutaminolysis, the other energy metabolic feature of tumor cells and proliferating normal cells, is also inefficient for energy production. Over consumption of glutamines by cancer cells is well documented, and it contributes to cachexia a main direct cause of death of cancer patients (*32*). However, in the higly proliferative glioblastoma cells, it was found that 60% of glutamine-derived carbons are not oxidized to produce ATP but rather secreted as lactates and alanines (*33*). Why the proliferating cells that have higher demand for energy than resting cells favor the low efficient Warburg effect and glutaminolysis for energy production is not fully understood. A prevalent view is that intermediates of these two pathways can be diverted to other pathways such as the pentose phosphate pathway (PPP) for biomass synthesis, and excess carbons are dumped in favor of the production of NADPH required for anabolic chemical reactions (*34*). However, anabolic pathways can also self regulated. For example, in highly proliferating cells reactions in the non-oxidative phase of PPP are reversed so that ribose-5-phosphate can be produced for nucleotide synthesis (*35*). In addition, intermediates of the TCA cycle, particularly citrate and oxaloacetate, can leave the cycle to support biomass synthesis (*8, 36*). Therefore, biomass synthesis is only partially dependent on areobic glycolysis and glutaminolysis. It is unclear whether such dependency is irreplaceable.

Here, we report evidence for an alternative explanation for the Warburg effect and glutaminolysis in proliferating cells. We show that influx of pyruvates or fatty acids to the TCA cycle lowers intracellular pH (pHi), which in turn induces apoptosis. In contrast, glutaminolysis and the Warburg effect increase pHi to allow the proliferating cells to survive.

## RESULTS

### Opposite effects of high and low pHi on cell proliferation

Normal lymph node cells were labeled with carboxyfluorescein succinimidyl ester (CFSE) and stimulated in vitro with the T cell mitogen anti-CD3 antibodies or the B cell mitogen LPS together with or without IL-2 or IL-4, respectively. Cell proliferation as measured by the serial dilution of CFSE was detected even in the absence of the mitogens, indicating the presence of pre-existing proliferating cells. (Fig. S1). Stimulation with the mitogens further increased the cell proliferation. Highly proliferating lymphocytes that had undergone 3 or more divisions were found primarily in cell populations with high pHi, nonetheless small numbers of the highly proliferating cells were also found in the cells with low pHi (Figs. 1A and S1). These results suggested that high pHi favored whereas low pHi was incompatible with the survival of the highly proliferating lymphocytes.

**Fig. 1.**
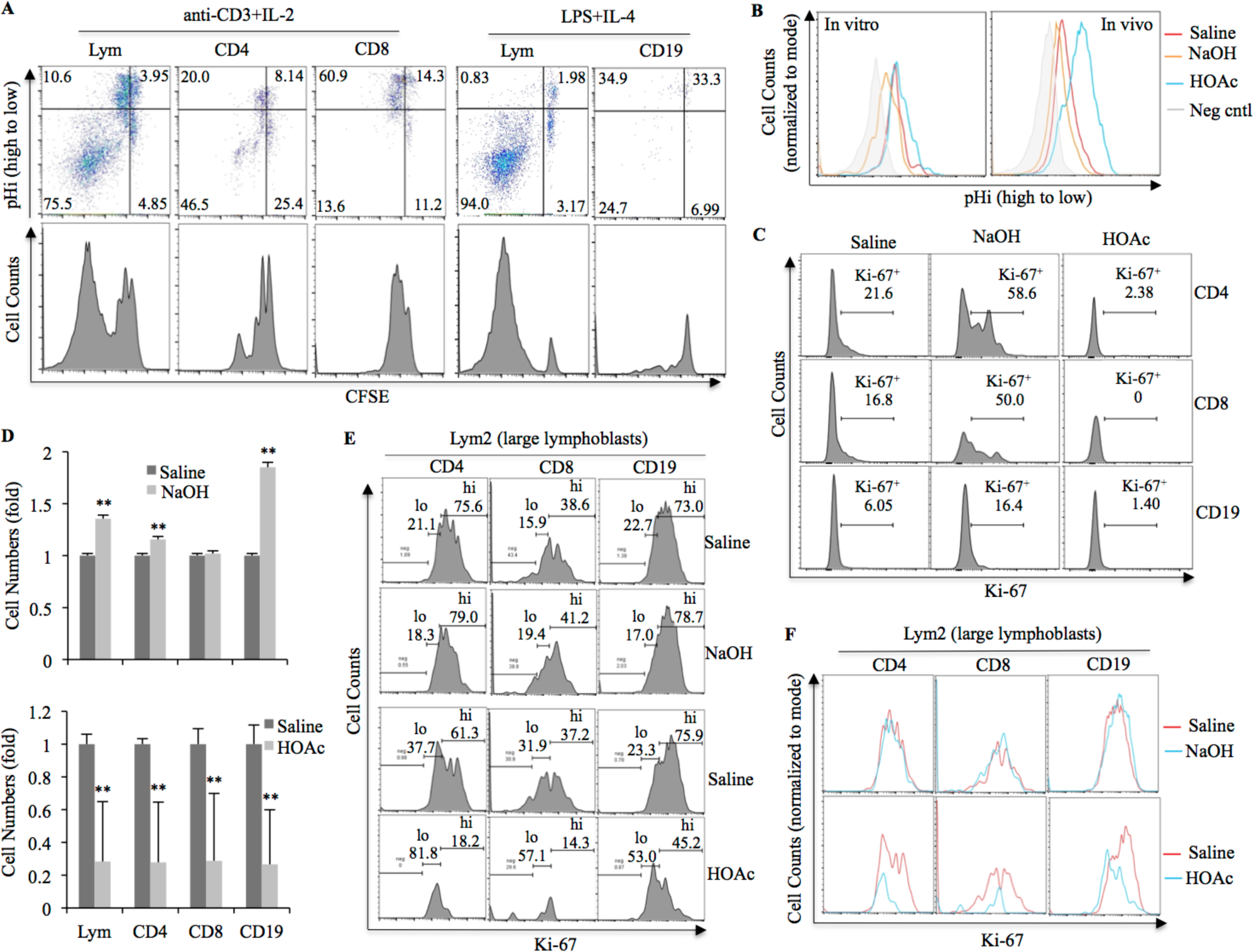
Regulation of lymphocyte proliferation by pHi. (A) CFSE-labeled lymph node cells were stimulated in vitro as indicated. Top panels show the staining of CFSE and the pHi indicator pHrodo^TM^ Red of live total lymphocytes (Lym), CD4 and CD8 T cells, and B cells (CD19). The lower panels are histograms of CFSE staining of the same cells. (B) Overlaid histograms of pHi of live total lymphocytes after in vitro or in vivo treatments with saline or saline plus NaOH or HOAc. Negative controls were live lymphocytes without pHi staining. (C) Histograms of Ki-67 staining in live CD4, CD8 T cells and B cells of lymph node cells treated in vitro. (D-F) Effects of in vivo treatments with NaOH or HOAc on lymphocytes in the MLNs of OVA-sensitized and challenged mice. (D) Average folds, relative to saline controls, of the numbers of live total lymphocytes, CD4 and CD8 T cells and B cells in mice treated with saline plus NaOH (upper) or HOAc (lower). Statistical significance of differences from saline controls was determined by Student *t* test, double asterisks indicate *P<0.01*. Pooled data of 3 or more similar experiments are shown. (E) Histograms of Ki-67 staining in the live Lym2 populations (large lymphoblasts) of the T and B cells. Numbers in the histograms are the percentages of Ki-67 high (hi) or low (lo) cells. (F) Overlaid histograms of Ki-67 staining of the same cells as in (E).

To further investigate the functional relation between pHi and lymphocyte proliferation, we sought to manipulate pHi by mild treatments with NaOH or HOAc solution prepared in saline. For in vitro treatments, lymphocytes were incubated in FBS to better mimic in vivo condition and avoid interference by pH buffering agents in regular cell culture medium. In vivo treatments were carried out by i.p. injection of the NaOH or HOAc solution. Both in vitro and in vivo, NaOH treatments effectively raised, whereas HOAc treatments lowered, the pHi in total peripheral lymphocytes (Fig. 1B), as well as in the CD4, CD8 T cells and B cells (Fig. S2).

The effects of the treatments on lymphocyte proliferation were studied by analyzing the level of Ki-67, a reliable marker for proliferating cells whose level positively correlates with rRNA and DNA synthesis (*37*). Consistent with the results in Fig. 1A, in vitro treatments with NaOH increased the percentages of Ki-67^+^ T and B cells and their levels of Ki-67 as compared with saline treatments. In contrast, in vitro HOAc treatments almost completely depleted the Ki-67^+^ but not the Ki-67^−^ lymphocytes. (Fig. 1C).

For the in vivo study, mice were sensitized and intratracheally challenged with ovalbumin (OVA). In the lung-draining lymph nodes (*38*), hereafter collectively referred to as mediastinal lymph nodes (MLNs), NaOH treatments modestly but significantly increased the number of live total lymphocytes, CD4 T cells and B cells (Fig. 1D, upper). In contrast, HOAc treatments dramatically reduced the number of all lymphocyte populations (Fig. 1D, lower). The MLNs contained lymphocytes of small and large sizes referred hereto as Lym1 and Lym2, respectively (Fig. S3A, D). The majority of the Lym1 cells expressed low levels of Ki-67, and NaOH treatments showed little effects on this population (Fig. S3G, left). Likewise, HOAc treatments showed only modest effects on Ki-67^lo^ CD4 T cells and B cells of the Lym1 populations, but caused sizable reduction of Ki-67^lo^ CD8 T cells (Fig. S3G, right). Unlike the Lym1 cells, the Lym2 cells contained large percentages of highly proliferating (Ki-67^hi^) cells (Fig. 1E). NaOH treatments increased the percentages of the Ki-67^hi^ cells and elevated Ki-67 levels in Ki-67^lo^ cells (Fig. 1E, F, upper). In contrast, HOAc treatments preferentially depleted the Ki-67^hi^ cells, and consequently increased the percentages of Ki-67^lo^ cells (Fig. 1E, F, lower).

### Induction of apoptosis by low pHi

We investigated whether the low pHi-induced death was mediated by apoptosis. Since pHi in apoptotic (dead) cells cannot be measured because of their leaky cell membranes, we instead examined early apoptosis of lymphocytes in the MLNs and non-draining lymph nodes (NDLNs). Early apoptotic (Annexin V^+^7-AAD^−^) cells were found mainly in the pHi^lo^ lymphocyte populations (Fig. 2A). Analysis of caspase-3 activation confirmed this finding, and demonstrated that the low pHi-induced apoptosis was mediated by caspase activation (Fig. 2B). To corroborate this finding, the MLN cells were treated with HOAc in vitro. The treatments caused dose-dependent induction of apoptosis in all lymphocyte populations (Fig. 2C), and again the early apoptotic cells were found primarily in the pHi^lo^ populations (Fig. 2D).

**Fig. 2.**
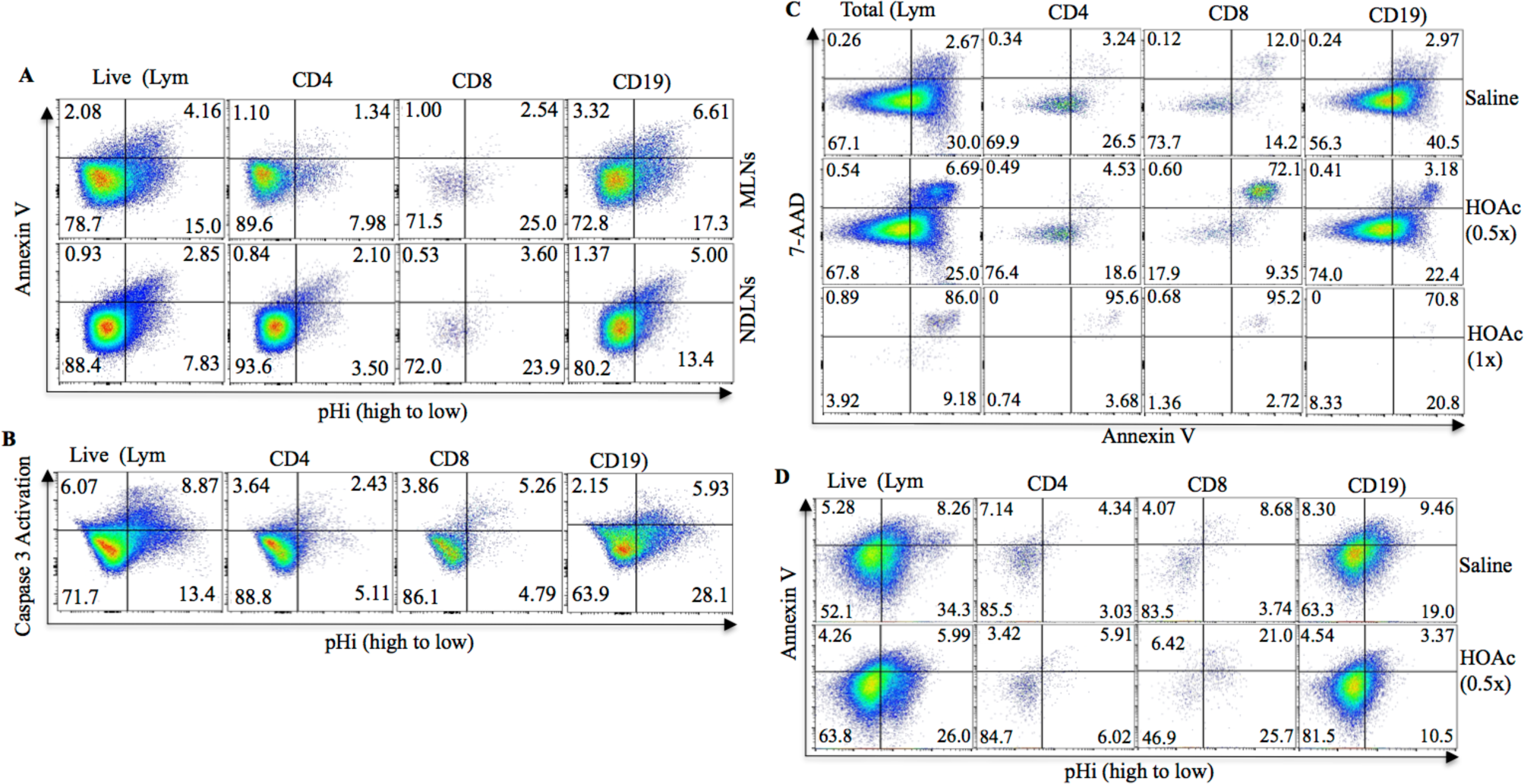
Induction of apoptosis by low pHi in MLN lymphocytes of OVA-sensitized and challenged mice. (A) Ex vivo lymphocytes from MLNs or NDLNs were stained with Annexin V and the pH indicator pHrodo^TM^ Green. Numbers in the plots are the percentages of cells in each quadrant. (B) Ex vivo lymphocytes stained with caspase-3 substrate NucView® 405 and pHrodo^TM^ Green. (C) MLN lymphocytes were incubated in FBS plus 1/10 volume of saline or saline containing 43.75mM (0.5x) or 87.5mM (1x) HOAc. Shown are plots of 7AAD and Annexin V staining of the indicated lymphocyte populations. (D) Annexin V and pHi staining of live (7AAD^−^) lymphocytes in (C).

### Inverse relation between MMPs and pHi at the early stage of immune response

To uncover the mechanisms for the regulation of pHi in lymphocytes, NDLNs and MLNs were harvested 3 days after the initial OVA challenge to measure the mitochondrial membrane potentials (MMPs) and pHi of the lymphocytes. Pseudocolor plots show that the vast majority of lymphocytes in the NDLNs displayed an inversion relation between MMPs and pHi, this relation was largely preserved, albeit less prominently, in lymphocytes in the MLNs (Fig. 3A, left). Comparisons between NDLNs and MLNs found that the pHi values of MLN lymphocytes were much higher than those of the NDLN lymphocytes whereas the opposite was found with the MMPs (Fig. 3B, right). We further divided the CD4, CD8 T cells and B cells in MLNs into distinct subpopulations of R1, R2, R3 and R4, which had progressively higher pHi (Fig. 3B, left). Scatter plotting of the pHi values of the subpopulations against their MMPs again showed an inverse relation between MMPs and pHi although this relation was relatively weak in the subpopulations with relatively high pHi (Fig. 3B, right).

**Fig. 3.**
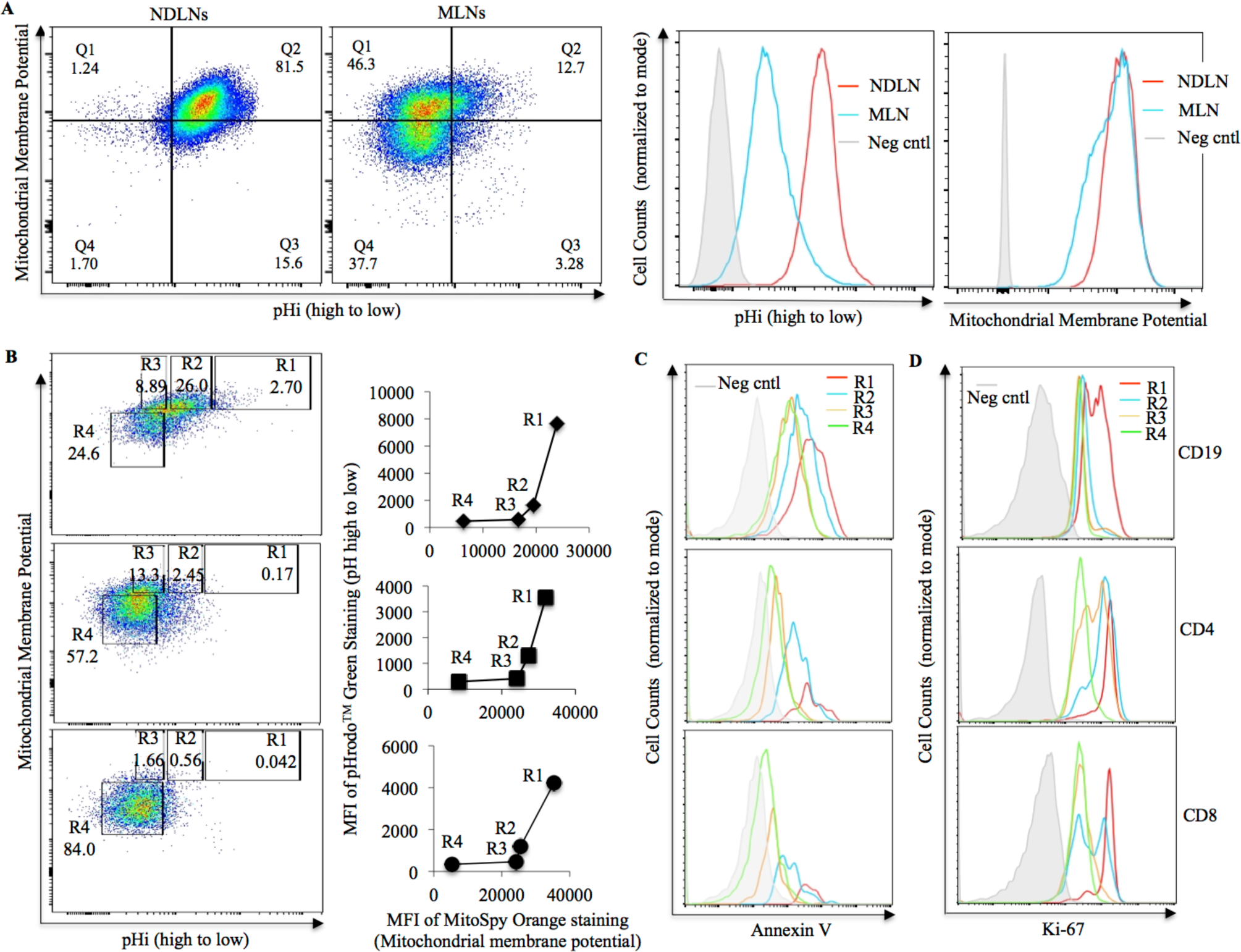
Interrelations among MMPs, pHi, apoptosis and cell proliferation at the early stage of immune response in the lymph nodes of OVA-sensitized and challenged mice. (A) Comparisons of MMPs and pHi of ex vivo live lymphocytes in the non-draining lymph nodes (NDLNs) and MLNs. Shown are pseudocolor plots of MMPs and pHi (left) and overlaid histograms of the pHi or MMP staining (right). (B) Inverse correlation between MMPs and pHi. Live T and B cells of the MLNs were divided into R1 to R4 subpopulations with progressively higher pHi (left). Shown on the right are scatter plots of pHi against MMPs of the different subpopulations. (C) Overlaid histograms of Annexin V staining of the R1 to R4 subpopulations. (D) Overlaid histograms of Ki-67 staining of sorted R1 to R4 subpopulations as defined in (B).

The functional roles of the pHi in apoptosis and cell proliferation were studied by comparing the Annexin V and Ki-67 staining of the R1 to R4 subpopulations. The highest intensities of Annexin V staining were detected in the R1 subpopulations that had the lowest pHi, the Annexin V staining decreased as the pHi increased (Fig. 3C). Similar patterns were observed with the Ki-67 staining (Fig. 3D). Notably, despite their high proliferative statuses, the R1 cells were of the lowest percentages (Fig. 3B, left), which is consistent with the in vitro finding that highly proliferating cells with low pHi failed to accumulate (Fig. 1A). Thus, at the early stage of the immune response, high rates of lymphocyte proliferation required high mitochondrial energetic activities, which however lowered pHi that in turn induced apoptosis.

### Dynamic changes of pHi and its relations with MMPs

We extended the study of MMPs and pHi to earlier and later time points. In these studies, the live total lymphocytes, CD4, CD8 T cells and B cells in the MLNs were divided into those with relatively high and low pHi referred hereto as the P and N lymphocytes, respectively. At the earlier time of day 2 after the initial OVA challenge, as well as in unimmunized mice, both the P and N lymphocytes showed inverse relation between MMPs and pHi. Nonetheless, there were higher percentages of P lymphocytes in the OVA-sensitized and challenged mice than in unimmunized mice (Fig. 4). At the intermediate stage of day 5, the percentages of P lymphocytes further increased, but there were still large percentages of N lymphocytes. More strikingly, at this stage, the P lymphocytes had assumed a positive relation between their MMPs and pHi, whereas the N lymphocytes maintained the inverse relation but were reduced in proportions (Fig. 4). As the immune response further progressed to the late stage, the P lymphocytes with a positive relation between MMPs and pHi became overwhelmingly the dominant populations, whereas the N lymphocytes with the inverse relation between MMPs and pHi were diminished to small minorities (Fig. 4).

**Fig. 4.**
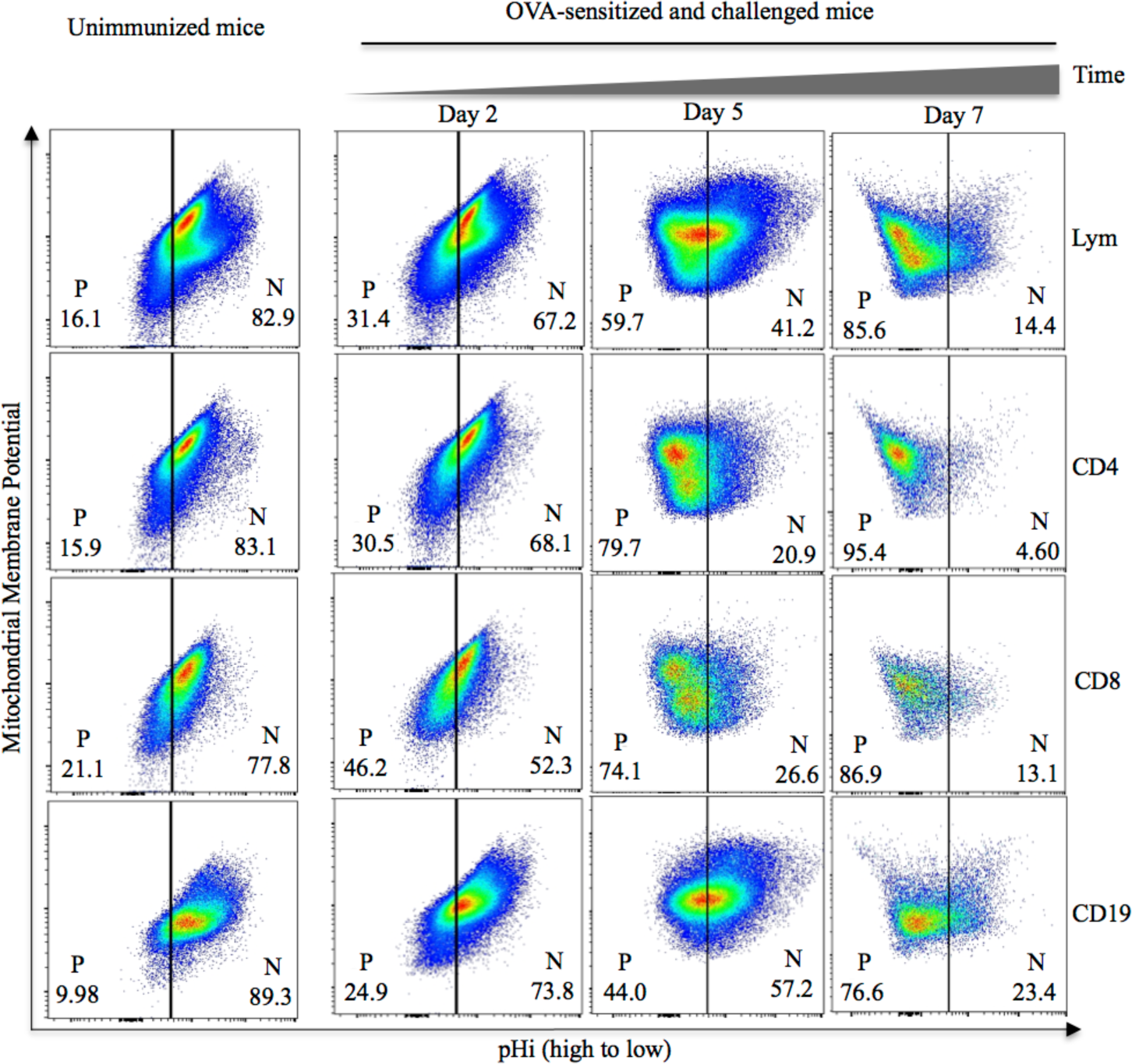
Dynamic changes of pHi and MMPs of lymphocytes in MLNs of OVA-sensitized and challenged mice. Shown are psuedocolor plots of MMP and pHi staining of ex vivo live lymphocytes. The lymphocytes are divided into populations of relatively high (P) and low (N) pHi. MLNs were harvested on day 2, 5 or 7 after the initial OVA challenge (day 0). Numbers in the plots are percentages of the N and P lymphocytes.

### Higher rates of proliferation but no apoptosis of MMPs-high P lymphocytes

MLN lymphocytes at the late stage of immune response were studied to determine the inter-relations among MMPs, pHi, apoptosis and proliferation in the P lymphocytes, and compare with the N lymphocytes. Like at the early stage of immune response in Fig. 3, early apoptotic cells were detected in the N lymphocytes, among which those with higher MMPs had higher percentages of early apoptotic cells than those with lower MMPs. In contrast, no distinct populations of early apoptotic cells were detected in the P lymphocytes regardless of their MMP levels (Fig. 5A, B). However, both the P and N lymphocytes with higher MMPs had higher rates of proliferation as judged by their Ki-67 levels than their counterparts with lower MMPs (Fig. 5C-E). Thus, unlike the N lymphocytes, P lymphocytes could proliferate, even at high rates, without triggering apoptosis.

**Fig. 5.**
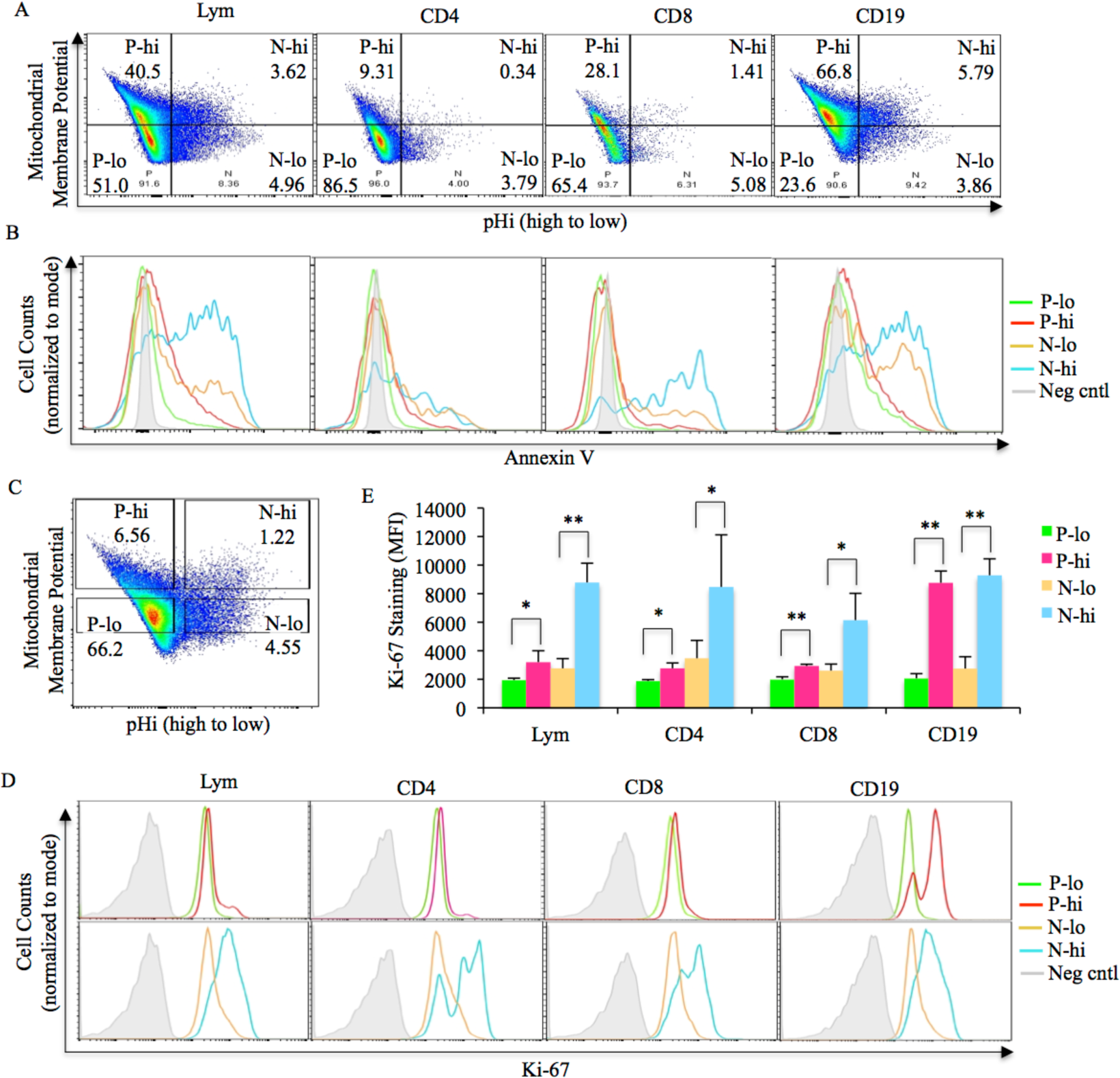
Interrelations among MMPs, pHi, apoptosis and proliferation of P lymphocytes and comparisons with N lymphocytes. (A) The P and N populations of T and B cells of MLNs are divided into subpopulations with low (lo) or high (hi) MMPs. (B) Overlaid histograms of Annexin V staining of the subpopulations defined in (A). (C-E) Ki-67 expression in the sorted subpopulations. (C) Sorting gates of the indicated subpopulations. (D) Overlaid histograms of Ki-67 staining of CD4, CD8 T cells or B cells of the sorted subpopulations. (E) Bar graph of the averages of mean fluorescence intensities (MFI) of Ki-67 staining of the sorted subpopulations. Positive error bars are standard deviations. Statistical significance of differences between the MMP-high and –low subpopulations of P and N lymphocytes, respectively, was determined by Student *t* test. Single asterisk indicates *P<0.05*, and double asterisks indicate *P<0.01*. Pooled data from 3 similar experiments are shown.

### Regulation of pHi and N and P lymphocyte proportions by energy metabolic pathways

The relations between MMPs and pHi suggested a role of energy metabolism in the regulation of pHi in lymphocytes and in the relative proportions of N and P lymphocytes. To test this, we used pharmacological compounds and MLN lymphocytes at the intermediate stage of immune response to pinpoint the energy metabolic pathways for such regulation. Compound GSK2837808A (GSK for short) is an inhibitor of lactate dehydrogenase therefore an inhibitor of the Warburg effect (*39*); CB-839 is an inhibitor of glutaminase 1 hence an inhibitor of glutaminolysis (*40*); dichloroacetate (DCA) is an inhibitor of pyruvate dehydrogenase kinase therefore promotes the influx of carbons derived from pyruvates to the TCA cycle (referred to hereafter as influx of pyruvates) (*41*); and C75 is an activator of carnitine palmitoyltransferase 1A therefore promotes the influx of carbons derived from fatty acids to the TCA cycle (referred to hereafter as influx of fatty acids) (*42*).

Inhibiting glutaminolysis or the Warburg effect, or alternatively increasing the influx of pyruvates or fatty acids, lowered the pHi of CD4, CD8 T cells and B cells (Fig. 6A, B). Therefore, gluatminolysis or the Warburg effect elevated whereas the influx of pyruvates or fatty acids lowered pHi. Consistent with this finding, inhibition of glutaminolysis and the Warburg effect reduced the percentages of the P lymphocytes (Fig. 6C, D) and their absolute numbers (Fig. 6E). Although the percentages of the N lymphocytes increased due to the reduction of the P lymphocytes, inhibition of either pathway decreased the absolute numbers of the N lymphocytes of the CD8 T cells and B cells, and did not significantly change the absolute numbers of the N lymphocytes of the CD4 T cells (Fig. S4). Although increase of the influx of pyruvates or fatty acids lowered pHi (Fig. 6A), it did not reduce the percentages of the P lymphocytes of the CD4 and CD8 T cells but it did reduce the percentages of the P lymphocytes of the B cells (Fig. 6C, D). However, regardless of the changes in the percentages of the P and N lymphocytes, the increase of influx of pyruvates or fatty acids reduced the absolute numbers of both the P and N lymphocytes of both T and B cells (Figs. 6E and S4). The reduction of the absolute cell numbers by inhibiting glutaminolysis or the Warburg effect, or alternatively by increasing the influx of pyruvates or fatty acids, is consistent with the notion that these interferences of energy metabolism lowered pHi therefore promoted apoptosis.

**Fig. 6.**
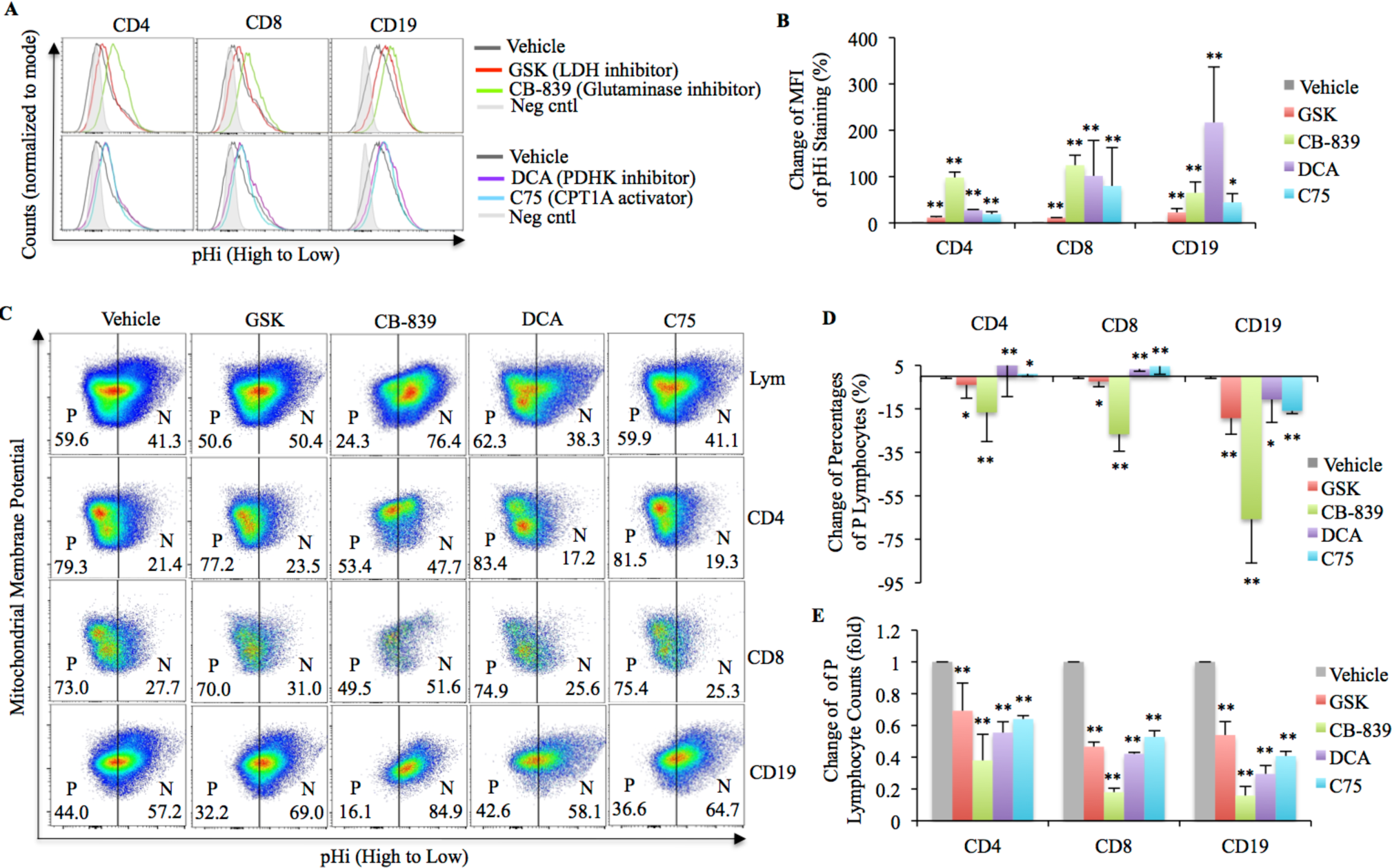
Regulation of pHi and the proportions of P and N lymphocytes by energy metabolic pathways. OVA-sensitized and challenged mice received i.p. injections of vehicle control or the indicated energy metabolic regulator. Ex vivo live lymphocytes of the MLNs from the mice were analyzed. (A) Overlaid histograms of pHrodo^TM^ Green staining of the T and B cells. (B) Bar graph of the average MFI of pHrodo^TM^ Green staining. (C) Pseudocolor plots of pHi and MMPs of live total lymphocytes (Lym), CD4, CD8 T cells and B cells. Numbers in the plots are the percentages of the P and N lymphocytes. (D) Bar graph showing the average changes of the percentages of P lymphocytes as compared with vehicle controls after the treatments. (E) Bar graph showing the average folds, relative to vehicle controls, of the cell counts of the P lymphocytes after treatments with the indicated energy metabolic regulators. Positive or negative error bars in all bar graphs are standard deviations. Statistical significance of differences from the vehicle controls was determined by Student *t* test. Single asterisk indicates *P<0.05,* double asterisks indicate *P<0.01*. Pooled data from 3 or more similar experiments are shown.

## DISCUSSION

While previous studies investigated the biological consequences of extracellular acidosis, in this study, we discovered a critical role of intracellular pH (pHi) in regulating the death and proliferation of lymphocytes hence the magnitude of immune response. We found that low pHi induces apoptosis, whereas high pHi promotes survival of proliferating lymphocytes. We further showed influx of carbons derived from pyruvates or fatty acids to the TCA cycle lowers pHi, whereas glutaminolysis and aerobic glycolysis raise pHi. Thus, the selective use of energy metabolic pathway determines the fate of proliferating lymphocytes. In other words, in addition to supporting biomass synthesis, glutaminolysis and the Warburg effect are indispensible survival strategy of the proliferating cells. Although our study focuses on lymphocytes, it is worth noting that glutaminolysis and the Warburg effect are found in various types of normal and malignant cells (*34, 43*). Moreover, the Warburg effect and the dependency on glutamines for cell proliferation are evolutionary conserved (*44–47*). Therefore, it is reasonable to speculate that the principles discovered in this study are applicable to other cell types across species, particularly cells that produce large amount of energy.

This study shows that low pHi is a natural and cell intrinsic trigger of lymphocyte apoptosis in both NDLNs and the lung-draining MLNs during an active immune response. This observation is distinct from cytosolic acidification in cultured cells as a result of the redistribution of mitochondrial contents caused by external agents that compromise the mitochondrial membrane integrity (*48–51*). In contrast, the low pHi observed in the lymphocytes in this study was at least in part the result of normal mitochondrial energy production fueled by carbons derived from pyruvates and fatty acids. On the other hand, we also observed a bi-phasic increase of pHi in lymphocytes. At the early stage of immune response, the increase of pHi is related to the reduction of MMPs. As the immune response progresses, the increase of pHi is largely dependent on glutaminolysis and aerobic glycolysis. Thus, this study identified energy metabolism as a major mechanism for regulating pHi in lymphocytes. In tumor cells, overactive acid extrusion and sequestration in organelles such as the Golgi apparatus lead to higher cytosolic pH than in untransformed cells (*52, 53*). Whether such mechanisms also play a part in the increase of pHi in the proliferating lymphocytes remains to be determined.

The dynamic nature of immune response stresses the importance of understanding and medically interfering with the molecular events in lymphocytes in accordance with the kinetics of immune response. However, unlike in laboratory settings, in clinical settings the time of initial antigen exposure is usually uncertain therefore it is hard to determine the stage in the natural time course of an immune response at the time of medical treatment. This study shows that the stage can be determined by examining the pHi of the lymphocytes and its relations with MMPs. Specifically, we found that lymphocytes can be divided into low and high pHi populations. The early stage of immune response is dominated by low pHi lymphocytes with an inverse relation between pHi and MMPs. As the immune response progresses to the intermediate stage, high pHi lymphocytes increase and assume a positive relation between pHi and MMPs. At the late stage, the immune response is dominated by high pHi lymphocytes with a positive relation between pHi and MMPs whereas the low pHi lymphocytes become small minorities. Thus, based on these characteristics, the stages of immune response can be determined without the knowledge of the time of antigen exposure. Application of this immune stage characterization system to clinical cases could improve the precision and efficacy of therapies targeting immune response.

## EXPERIMENTAL PROCEDURES

### Mice and OVA immunization model

Balb/c mice were purchased from Jackson Laboratory (Bar Harbor, ME) and housed in the animal facility of Charles River Accelerator and Development Lab (CRADL) (Cambridge, MA). Additional mice were maintained at Therazwimm animal facility. Animal studies were performed according to the protocols approved by the CRADL and Therazwimm Institutional Animal Care and Use Committees. OVA sensitization and challenge were performed as previously described (*54, 55*). Briefly, Balb/c mice were sensitized by i.p. injection of 20μg OVA (Sigma-Aldrich, St. Louis, MO) mixed with Alum adjuvant (Thermo Scientific, Rockford, IL). Two weeks later, the sensitization is repeated with 500μg OVA mixed in Alum adjuvant. Two weeks later, mice were challenged with 100μg OVA in 60μl saline by intratracheal (i.t.) instillation. The challenge was repeated as indicated in other procedures.

### In vitro stimulation of lymphocytes

Lymph node cells were washed 3 times with plain PBS, then incubated (5 x 10^6^/ml) in 1μm Carboxyfluorescein succinimidyl ester (CFSE) (Fluka/Sigma-Aldrich, Burlington, VT) in plain PBS at room temperature for 7 minutes. After the incubation, 1/4 volume of FBS was added to stop the labeling, and cells were washed 4 times with PBS plus 1% FBS. The labeled cells were cultured in complete RPMI-1640 medium plus IL-2 (20 units/ml) or IL-2 and anti-CD3 antibody (1μg/ml) (BD Pharmingen (San Diego, CA), or alternatively plus IL-4 (4ng/ml) or IL-4 and lipopolysaccharide (LPS) (10μg/ml) (Sigma-Aldrich, St. Louis, MO) for 2.5 days.

### Measuring pHi, MMP and early apoptosis

MMP and pHi were determined by staining lymphocytes with the MMP indicator MitoSpy Orange (Biolegend, San Francisco, CA) and the pH indicator pHrodo^TM^ Green/Red AM (ThermoFisher Scientific, Waltham, MA), respectively. Intensity of MitoSpy staining is MMP-dependent, and intensity of pHrodo^TM^ staining is inversely correlated with pHi value. Apoptosis and early apoptosis were detected by staining the lymphocytes with 7AAD and Annexin V (Biolegend) or NucView® Caspase-3 Substrate (Biotium, Fremont, CA). For staining with MitoSpy, pHrodo^TM^ and Annexin V, cells were washed once with Live Cell Image Solution (LCIS) (Life Technoology/ThermoFisher Scientific, Grand Island, NY). The cells (1-2 x 10^7^/ml) were re-suspended in LCIS supplemented with 0.7mM CaCl_2_, plus 1μM pHrodo^TM^, 100nM MitoSpy Orange, Annexin V and/or the Caspase-3 Substrate, and Zombie viability dye (Biolegend) at dilutions recommended by the manufacturers individually or in combination. The cell suspension was incubated in 37°C water bath for 20 minutes and in ice water for 15 minutes, then washed in LCIS supplemented with 0.7mM CaCl_2_ and 1% FBS. The cells were re-suspended in LCIS supplemented with 0.7mM CaCl_2_, 7-AAD was added to the cells prior to flow cytometric analysis. MMP, pHi and early apoptosis were determined in gated live (7AAD^−^ or Zombie^−/lo^) lymphocytes.

### Flow cytometry

Fluorochrome-conjugated antibodies against mouse CD4, CD8, CD19, TCRβ and Ki-67, 7-AAD, Zombie-Green/Violet fixable viability dyes and Foxp3 buffer set were purchased from Biolegend (San Diego, CA). Lymphocytes were stained for cell surface markers. Live and dead cells were distinguished by staining the cells with the Zombie dyes or 7-AAD. For Ki-67 staining, cells were fixed with CytoFix/CytoPerm buffer, washed with PBS plus 1% FBS and 1x CytoPerm buffer, and incubated with antibodies against mouse Ki-67. For lymphocyte cell count by flow cytometry, lymph node cells of different mice were prepared as single cell suspensions in the same volume of buffer. An aliquot of the same volume of cell suspension were stained with 7AAD with or without staining with other dyes and antibodies. The stained cells were re-suspended in the same volume of buffer and analyzed by flow cytometry with the same acquisition volume, flow speed and acquisition time. All flow cytometry experiments were carried out using FACSCanto (BD. Bioscience, Franklin Lakes, NJ) or Attune^TM^ cytometer (Invitrogen, Carlsbad, CA).

### Treatments with pH modifiers or energy metabolic regulators

Lymphocytes were treated with NaOH or HOAc to determine the effects of the pH modifiers on pHi and cell proliferation. For in vitro treatments, lymphocytes were incubated in FBS containing 10% volume of saline or saline plus 87.5mM NaOH or HOAc in 37°C water bath for 5 hours for cell proliferation study and 20 minutes for determining effects on pHi. For in vivo treatments with NaOH, OVA-sensitized mice were challenged with OVA every other day for a total of three challenges, mice received i.p. injection of 200μl saline or saline plus 87.5mM NaOH 1 hour after each challenge, mice were sacrificed on day 3 after the final challenge. For in vivo treatments with HOAc, OVA-sensitized mice were challenged on day 0 and day 1, mice receive i.t. instillation of 60μl saline or saline plus 175mM HOAc on day 2, the mice were sacrificed on day 3. For in vivo treatments with energy metabolic regulators, OVA-sensitized mice were challenged on day 0 and day 1, and on day 2 and day 3 received i.p. injection of 200μl vehicle (saline plus 5% DMSO) or vehicle plus 48nM GSK2837808A, 10μM CB-839, 10mM dichloroacetate, or 1mM C75, mice were sacrificed 1 day (on day 5) after the second treatments. All energy metabolic regulators were purchased from MedChemExpress (Monmouth Junction, NJ).

## Acknowledgments

WZ thanks Doug Green for suggesting measuring caspase-3 activation to detect early apoptosis.

## Author contributions

WZ conceived, designed and carried out the study, and wrote the manuscript. SY and BZ assisted with flow cytometry data acquisition.

## Competing interests

WZ is the inventor of patent applications based in part on data in this article.

## Data and materials availability

All data are available in the main text or the supplementary materials.

## Supplementary Materials

Figs. S1 to S4

## Supplementary figures

**Figure S1.**
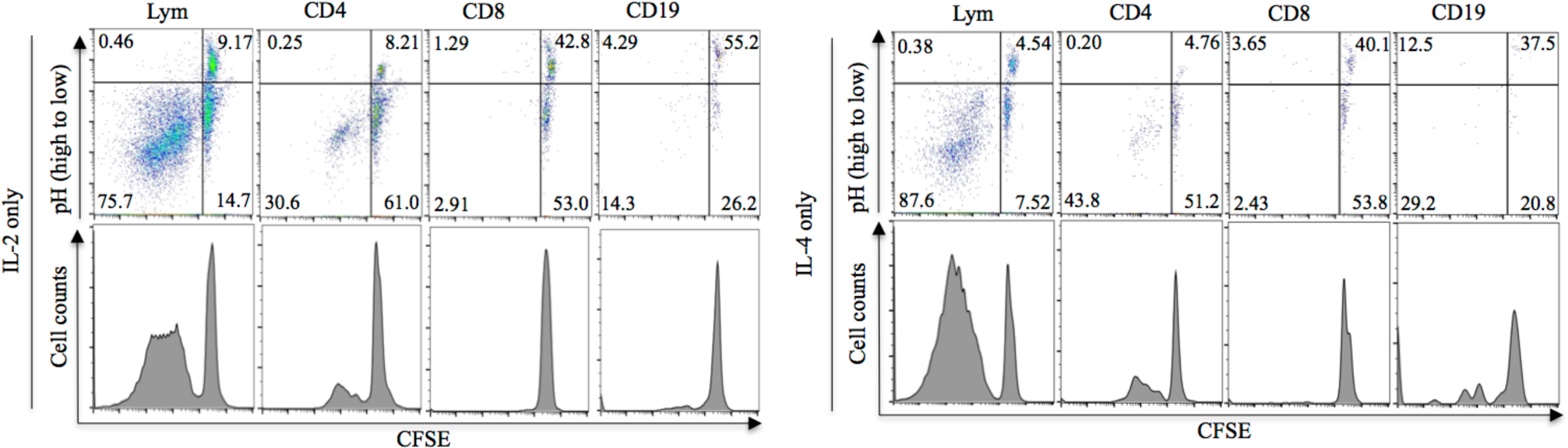
Proliferation of lymphocytes cultured in vitro with IL-2 (left) or IL-4 (right). Top panels are pseudocolor plots of CFSE staining and pHi of live total lymphocytes (Lym), CD4, CD8 T cells, and B cells (CD19^+^). The lower panels are histograms of CFSE staining of the same cells. Numbers in the plots are percentages of cells in their respective populations.

**Figure S2.**
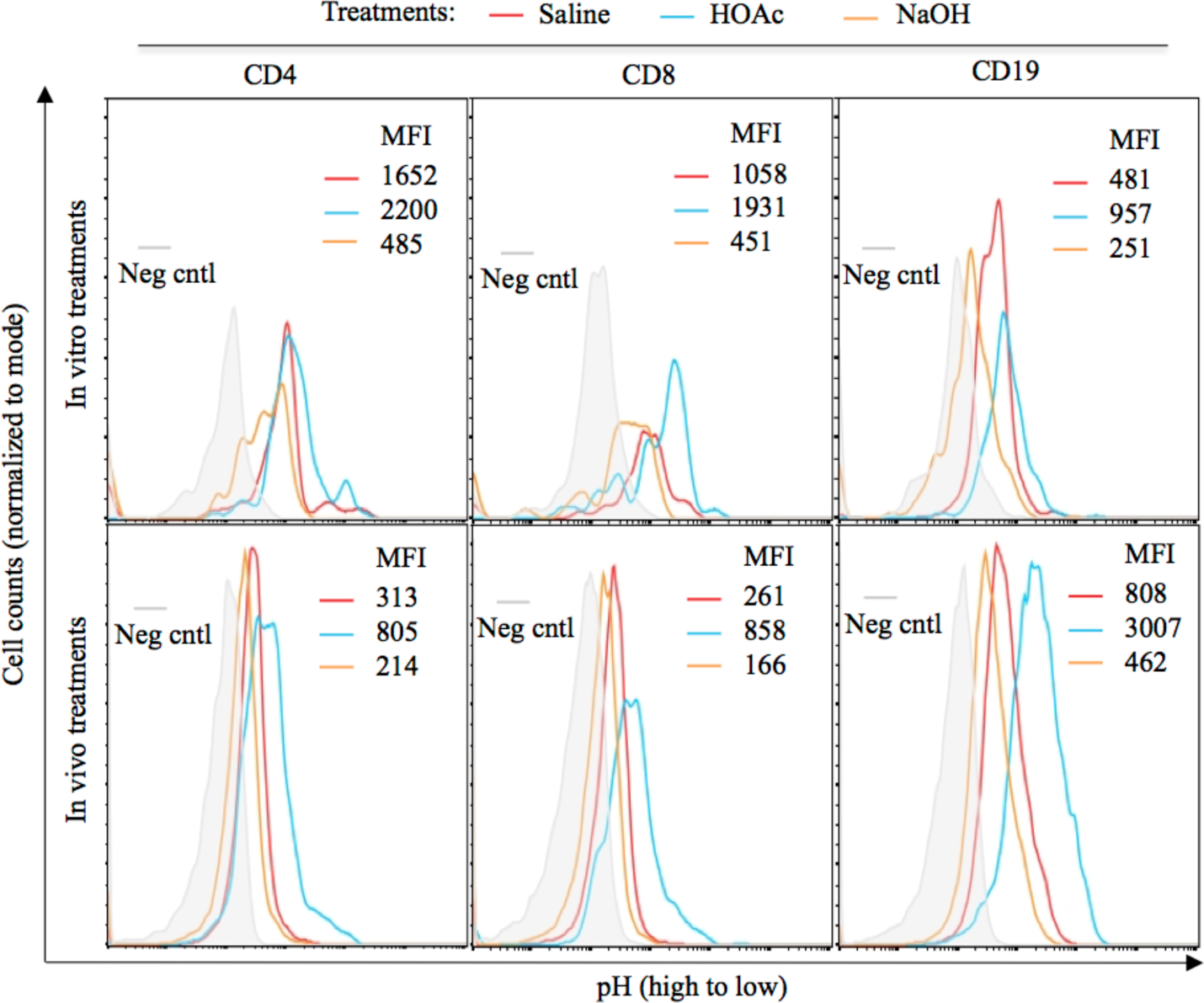
Intracellular pH in CD4, CD8 T cells and B cells (CD19) after in vitro (upper) or in vivo (lower) treatments with saline, saline plus HOAc or NaOH. Shown are the overlaid histograms and mean fluorescence intensities (MFI) of pHrodo^TM^ Green staining.

**Figure S3.**
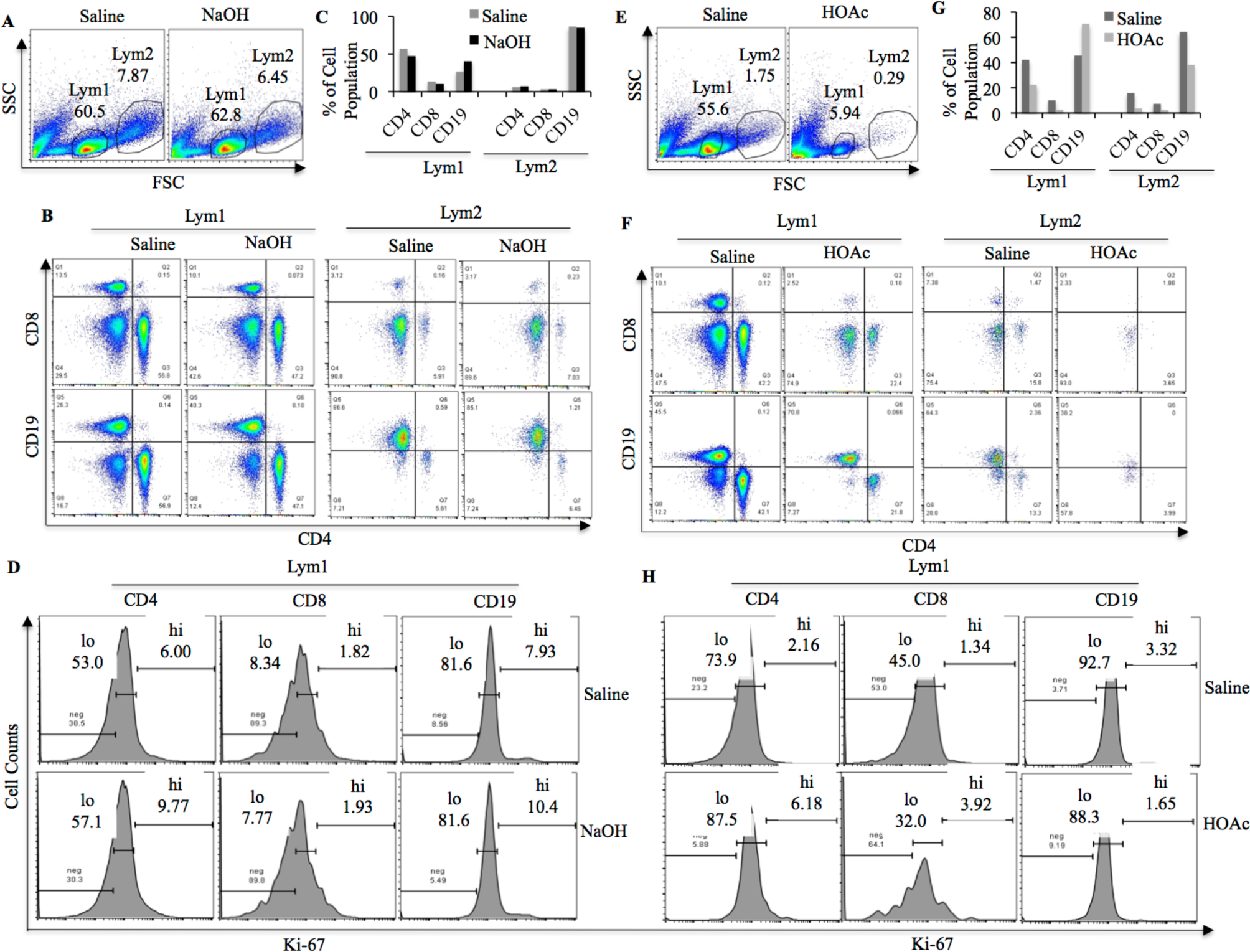
Lymphocyte populations in the MLNs of OVA-sensitized and challenged mice treated with saline or saline plus NaOH (A-D); or alternatively with saline or saline plus HOAc (E-H). (A, E) Pseudocolor plots of live (Zombie-Green^lo/-^) single cells (and some cell debris that are Zombie-Green^−^), showing the relatively small (Lym1) and large (Lym2) lymphocyte populations. (B, F) Pseudocolor plots showing the CD4, CD8 T cells and B cells (CD19^+^) populations in Lym1 and Lym2. (C, G) Bar graphs showing the percentages of CD4, CD8 T cells and B cells in (B) and (F), respectively. (D, H) Histograms of Ki-67 staining of CD4, CD8 T cells and B cells in the Lym1 populations. Numbers are the percentages of Ki-67-high (hi) or –low (lo) cells.

**Figure S4.**
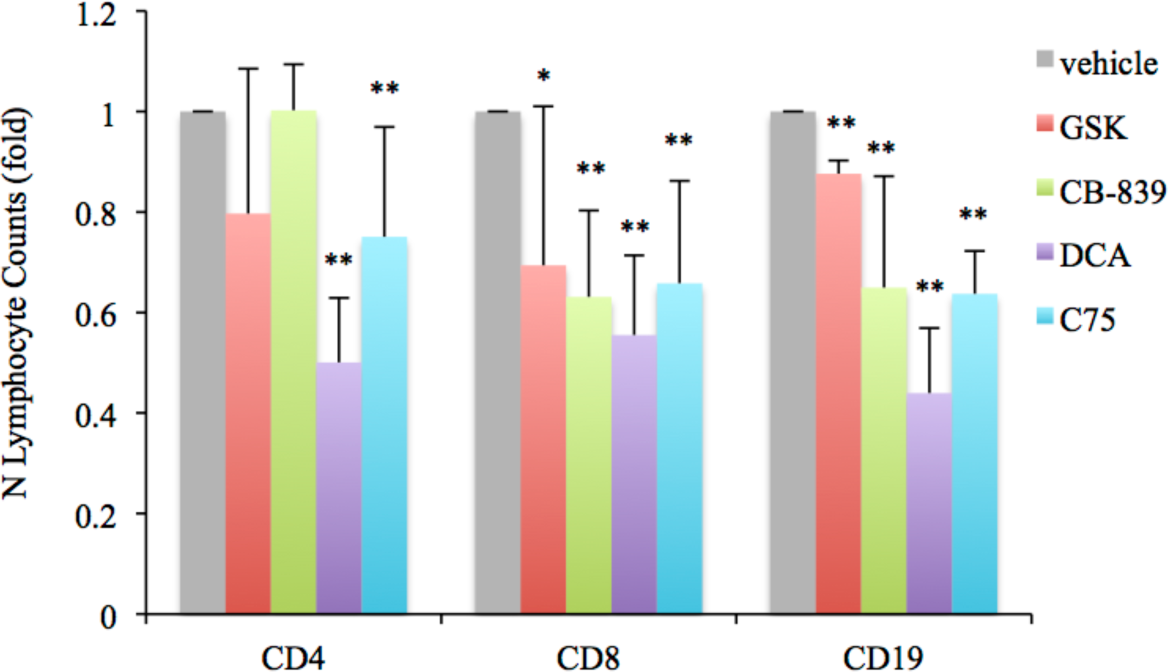
Bar graph showing the average folds, relative to vehicle controls, of the cell counts of the N lymphocytes after treatments of OVA-sensitized and challenged mice with the indicated energy metabolic regulators. Positive error bars are standard deviations. Statistical significance of differences from the vehicle controls was determined by Student *t* test. Single asterisk indicates *P<0.05,* double asterisks indicate *P<0.01*.

## Notes

### Summary of Updates

More data are included and title is changed.

